# Medical consequences of pathogenic CNVs in adults: Analysis of the UK Biobank

**DOI:** 10.1101/264077

**Authors:** Karen Crawford, Matthew Bracher-Smith, Kimberley M Kendall, Elliott Rees, Antonio F Pardiñas, Mark Einon, Valentina Escott-Price, James TR Walters, Michael C O’Donovan, Michael J Owen, George Kirov

**Affiliations:** MRC Centre for Neuropsychiatric Genetics & Genomics, Cardiff University, School of Medicine, Hadyn Ellis Building, Maindy Road, Cardiff CF24 4HQ, UK

## Abstract

**Background:** Genomic copy number variants (CNVs) increase risk for early-onset neurodevelopmental disorders but their impact on medical outcomes in later life is poorly understood. The UK Biobank, with half a million well-phenotyped adults, presents an opportunity to study the medical consequences of CNV in middle and old age.

**Methods:** We called 54 CNVs associated with clinical phenotypes or genomic disorders, including their reciprocal deletions or duplications, in all Biobank participants. We used logistic regression analysis to test CNVs for associations with 58 common medical phenotypes.

**Findings:** CNV carriers had an increased risk of developing 37 of the 58 phenotypes at nominal levels of statistical significance, with 19 associations surviving Bonferroni correction (p<8·6×10^−4^). Tests of each of the 54 CNVs for association with each of the 58 phenotypes identified 18 associations that survived Bonferroni correction (p<1·6×10^−5^) and a further 57 that were associated at a false discovery rate (FDR) threshold of 0·1. Thirteen CNVs had three or more significant associations at FDR=0·1, with the largest number of phenotypes (N=15) found for deletions at 16p11·2. The most common CNVs (frequency 0·5-0·7%) have no or minimal impact on medical outcomes in adults.

**Interpretation:** Some of the 54 tested CNVs have profound effects on physical health, even in people who have largely escaped early neurodevelopmental outcomes. Our work provides clinicians with a morbidity map of potential outcomes among carriers of these CNVs.

**Funding:** MRC UK, Wellcome Trust UK

## Background

Genomic copy number variations (CNVs) are structural alterations to chromosomes of >1,000 bases in length and can intersect multiple genes.^1^ CNVs at specific loci have been shown to increase risk for autism spectrum disorders,^2^ developmental delay and other neurodevelopmental disorders,^3^ and schizophrenia.^4^ Apart from their association with neurodevelopmental and psychiatric outcomes, these CNVs can lead to medical disorders. Several CNVs, for example deletions at 22q11·2,^5^ have been extensively studied on hundreds of carriers and their medical consequences are well established. However, for CNVs with lower penetrance, very rare CNVs or several reciprocal deletions/duplications of known genomic disorders, the associated medical phenotypes have not been identified. Moreover, most research has been performed on children and young people referred to genetic clinics,^3,6^ creating a referral bias towards early-onset medical conditions and more adverse outcomes. Most CNVs display incomplete penetrance for neurodevelopmental and congenital disorders,^7^ but the question whether apparently unaffected carriers experience an increased rate of medical outcomes in later life has not been addressed in adequately powered studies to date.

The establishment of the UK Biobank presents a unique opportunity to examine the spectrum of medical outcomes of CNVs in middle and old-aged people, as all half a million participants have been assessed with identical methods and blindly to their CNV status. The Biobank collects longitudinal data from hospital admissions, self-report, death certificates, cancer registries and primary care records. Here, we report on the medical consequences of carrier status for 54 well-recognised CNVs, reported to be associated with clinical phenotypes or genomic disorders,^3,6,8^ including their reciprocal deletions/duplications.

## Methods

### Participants

The UK Biobank recruited just over half a million people from the general population of the UK. Participants were between 40 and 69 years of age at the time of recruitment between 2006 and 2010. They consented to provide personal and health information, urine, saliva and blood samples, including for DNA testing. We report on the 421,274 participants who passed genotyping QC filters (see Methods) and were of white British or Irish descent. 54·1% of them were female and the mean age at the end of the current follow-up interval for medical outcomes (in 2016) was 64·7 years, SD=8·0 years.

### CNV calling

We obtained permission from the UK Biobank to analyse the CNVs in project 14421 : “Identifying the spectrum of biomedical traits in adults with pathogenic copy number variants (CNVs)”. The anonymised genotypic data from Affymetrix Axiom or BiLEVE arrays were downloaded as 488,415 raw (CEL) files from the UK Biobank, and analysed with the methods reported previously.^9^ Briefly, we generated normalised signal intensity data, genotype calls and confidences, using ~750,000 biallelic markers. These were then processed with PennCNV-Affy software.^10^ Individual samples were excluded if they had >30 CNVs, a waviness factor >0·03 or <-0·03, a call rate <96%, or Log R Ratio standard deviation >0·35.

A total of 25,069 files were excluded after this QC. Individual CNVs were excluded if they were covered by <10 probes or had a density coverage of less than one probe per 20,000 base pairs.

### Choice of CNVs

We compiled a list of 92 CNVs in 47 genomic locations from two widely accepted sources that proposed largely overlapping sets of CNVs (Table S1).^3,6^ The authors of these studies used information from databases, reviews and publications to produce lists of CNV regions that lead to genomic disorders, neurodevelopmental or other clinical phenotypes and congenital malformations. We refer to this set of 92 CNVs as “pathogenic”, consistent with the criteria proposed by the American College of Medical Genetics standards which describe as pathogenic those CNVs that have been documented as clinically significant in multiple peer-reviewed publications, even if penetrance and expressivity of the CNV are known to be variable.^11^ Many (but not all of these CNVs) have been shown to statistically increase risk for developmental delay. Table S1 lists the sources for selection and our criteria for inclusion in analysis. Several overlapping or adjacent CNVs listed as different loci in the original publications were grouped together (e.g. the “small” and the “common” 22q11·2, or the “small” and the “large” 16p13·3 deletions/duplications). As a rule, the reciprocal deletions/duplications of known genomic disorders were also included by the above authors and by us, in order to examine their medical consequences, even if the evidence for their pathogenicity has not been established.

The criteria for calling CNVs that do not span the full critical region are detailed in our previous work.^9^ As a rule, a CNV had to intersect at least 50% of the critical region (marked as “location” in Table 1 and Table S1) and affect the candidate genes which are thought to be relevant for pathogenesis, if known. For single gene CNVs, we required deletions to intersect at least one exon, and duplications to span the whole coding region. We observed several loci, mostly telomeric, where a number of small CNVs were preferentially called on arrays that failed QC (marked “Unreliable” in Table S1). In order to avoid potential false-positives on this genotyping platform, we excluded these loci from analysis. We also excluded from analysis CNVs with fewer than five observations in the full sample, as being too rare for analysis (marked “Rare” in Table S1). The above filtering left 54 CNVs for analysis (Table 1 and Table S1).

**Table 1.** List of CNVs analysed in the current study. “N results p<0-05” shows the number of nominally significant results from associations between the CNV and 58 medical phenotypes. The last column shows the number of Benjamini-Hochberg significant results at FDR=0·1 (“B-H FDR=0·1”).

| **CNV locus** | **Location (hg19)** | **N carriers** | **N results p<0.05** | **B-H FDR=0.1** |
| --- | --- | --- | --- | --- |
| TAR_del | chr1:145,394,955-145,807,817 | 75 | 7 | 0 |
| TAR_dup | chr1:145,394,955-145,807,817 | 436 | 10 | 1 |
| 1q21·1del | chr1:146,527,987-147,394,444 | 113 | 8 | 3 |
| 1q21·1dup | chr1:146,527,987-147,394,444 | 177 | 14 | 3 |
| NRXN1del | chr2:50,145,643-51,259,674 | 163 | 3 | 1 |
| 2q11·2del | chr2:96,742,409-97,677,516 | 31 | 2 | 0 |
| 2q11·2dup | chr2:96,742,409-97,677,516 | 29 | 2 | 0 |
| 2q13del(*NPH1*) | chr2:110,862,716-110,983,948 | 2448 | 6 | 1 |
| 2q13dup(*NPH1*) | chr2:110,862,716-110,983,948 | 1976 | 0 | 0 |
| 2q13del | chr2:111,394,040-112,012,649 | 53 | 14 | 1 |
| 2q13dup | chr2:111,394,040-112,012,649 | 71 | 3 | 1 |
| 2q21·1del | chr2:131,481,308-131,930,677 | 41 | 3 | 0 |
| 2q21·1dup | chr2:131,481,308-131,930,677 | 59 | 3 | 0 |
| 3q29del | chr3:195,720,167-197,354,826 | 9 | 7 | 1 |
| 3q29dup | chr3:195,720,167-197,354,826 | 5 | 14 | 6 |
| WBS_dup | chr7:72,744,915-74,142,892 | 14 | 0 | 0 |
| 7q11·23dup_distal | chr7:75,138,294-76,064,412 | 24 | 4 | 0 |
| 8p23·1dup | chr8:8,098,990-11,872,558 | 6 | 3 | 0 |
| 10q11·21q11·23del | chr10:49,390,199-51,058,796 | 57 | 4 | 0 |
| 10q11·21q11·23dup | chr10:49,390,199-51,058,796 | 43 | 8 | 0 |
| 10q23dup | chr10:82,045,472-88,931,651 | 7 | 6 | 0 |
| 13q12del(*CRY1*) | chr13:20,977,806-21,100,012 | 379 | 3 | 0 |
| 13q12dup(*CRY1*) | chr13:20,977,806-21,100,012 | 10 | 5 | 0 |
| 13q12·12del | chr13:23,555,358-24,884,622 | 85 | 6 | 0 |
| 13q12·12dup | chr13:23,555,358-24,884,622 | 236 | 1 | 0 |
| 15q11·2del | chr15:22,805,313-23,094,530 | 1664 | 3 | 0 |
| 15q11·2dup | chr15:22,805,313-23,094,530 | 2041 | 6 | 0 |
| PWS_dup | chr15:22,805,313-28,390,339 | 19 | 8 | 0 |
| 15q11q13del_BP3-BP4(*APBA2*, *TJP*) | chr15:29,161,368-30,375,967 | 16 | 10 | 1 |
| 15q11q13dup_BP3-BP4(*APBA2*, *TJP*) | chr15:29,161,368-30,375,967 | 53 | 2 | 0 |
| 15q11q13dup_BP3-BP5 | chr15:29,161,368-32,462,776 | 9 | 3 | 0 |
| 15q13·3del | chr15:31,080,645-32,462,776 | 42 | 12 | 3 |
| 15q13·3dup | chr15:31,080,645-32,462,776 | 240 | 10 | 2 |
| 15q13·3del(*CHRNA7*) | chr15:32,017,070-32,453,068 | 10 | 5 | 0 |
| 15q13·3dup(*CHRNA7*) | chr15:32,017,070-32,453,068 | 3031 | 2 | 0 |
| 15q24dup | chr15:72,900,171-78,151,253 | 9 | 7 | 0 |
| 16p13·11del | chr16:15,511,655-16,293,689 | 131 | 9 | 1 |
| 16p13·11dup | chr16:15,511,655-16,293,689 | 828 | 6 | 3 |
| 16p12·1del | chr16:21,950,135-22,431,889 | 246 | 19 | 8 |
| 16p12·1dup | chr16:21,950,135-22,431,889 | 202 | 3 | 0 |
| 16p11·2distal_del | chr16:28,823,196-29,046,783 | 58 | 7 | 4 |
| 16p11·2distal_dup | chr16:28,823,196-29,046,783 | 137 | 7 | 1 |
| 16p11·2del | chr16:29,650,840-30,200,773 | 110 | 28 | 15 |
| 16p11·2dup | chr16:29,650,840-30,200,773 | 138 | 8 | 2 |
| 17p12del(HNPP) | chr17:14,141,387-15,426,961 | 237 | 6 | 1 |
| 17p12dup(CMT1A) | chr17:14,141,387-15,426,961 | 124 | 12 | 4 |
| Potocki-Lupski Syndrome | chr17:16,812,771-20,211,017 | 5 | 7 | 0 |
| 17q11·2del(*NF1*) | chr17:29,107,491-30,265,075 | 9 | 5 | 0 |
| 17q12del | chr17:34,815,904-36,217,432 | 9 | 13 | 3 |
| 17q12dup | chr17:34,815,904-36,217,432 | 101 | 6 | 1 |
| 22q11·2del | chr22:19,037,332-21,466,726 | 10 | 7 | 0 |
| 22q11·2dup | chr22:19,037,332-21,466,726 | 280 | 20 | 4 |
| 22q11·2distal_del | chr22:21,920,127-23,653,646 | 5 | 12 | 3 |
| 22q11·2distal_dup | chr22:21,920,127-23,653,646 | 13 | 4 | 0 |

### Choice of medical phenotypes

Data on health outcomes were collected from several sources. Self-declared illnesses were disclosed by participants at their initial assessments and coded into 445 distinct categories. Hospital discharge diagnoses (primary and secondary) and death certificates contain over 11,000 ICD10 codes assigned to at least one participant. Analysing each individual code separately against 54 CNV loci would result in small numbers of participants with each code and thus not provide sufficient statistical power to detect the true associations. To reduce the dimentionality of the data and therefore increase power, we grouped together disease entities into broader disease groups, using all available sources. We gave preference to common conditions and grouped disorders into recognised categories, based on organ, system, or aetiology, while excluding from the current analysis infectious diseases, injuries, and neuropsychiatric disorders. The disease codes used to construct each phenotype group are listed in Table S2. For myocardial infarction and stroke we used the “adjudicated” data provided by the UK Biobank (datafields 42000 to 42013). Phenotype groups found in fewer than 2000 participants were not included. The final list of disease groups includes 58 entities, including “death during follow-up” obtained from the death registries (Table 1). Data on cancer were taken only from the UK cancer registries, as collected and supplied by the UK Biobank, as this is the most reliable and complete resource. For the current work we considered all malignant cancers as a single phenotype. As risk for cancer was not significantly affected by CNVs, and because most individual cancers affected relatively small numbers of patients, we did not analyse the cancers by sub-type.

### Statistical analysis

Analyses were performed in the statistical package R (version 3·3·0). We examined the effect of the presence of a CNV on the presence of a phenotype in logistic regression analysis with age and gender as co-variates. We used Firth's bias-reduced logistic regression method,^12^ with the library “logistf”, as it better handles cells with small numbers. We report the resulting p-value, odds ratio (OR) and 95% confidence intervals for the OR. We also report the Relative Risk (RR), for having the phenotype in carriers of a specific CNV and noncarriers of any of the 54 CNVs.

As the lifetime prevalence of disorders often varies by ancestry, we restricted the analysis to those participants who declared themselves as “white British or Irish”, a total of 421,274 participants with good quality CNV calls. Conservative Bonferroni correction was applied for the testing of 54 CNVs × 58 phenotypes, giving a p<1·6×10^−5^ as a project-wide significance level. We also report the Benjamini-Hochberg correction for a false discovery rate (B-H FDR) of 0·1.^13^ Table S3 shows all 3132 CNV/phenotype comparisons, grouped by CNV, including all corrected and uncorrected p-values. These data and additional figures for every CNV and every phenotype association are available at http://kirov.psycm.cf.ac.uk/. Table S4 displays the top Benjamini-Hochberg p-values, in ranked order down to p<0·2 (FDR=0·2).

## Findings

### CNV carriers have an increased rate of medical phenotypes

We first analysed how the chosen CNVs, as a group, affected the 58 medical phenotypes. For this group analysis we excluded the five relatively common CNVs (found in over 1500 carriers each, Table 1 and S1). One (or more) of these five CNVs was found in 11,049 people, while the remaining 49 CNVs were found in a total of 5129 people. Inclusion of the five common CNVs would effectively represent a test of the effect of these five CNVs, as they are present in over two thirds of carriers. (We do, however, provide all results in Table S3 and the website). Over half of the medical phenotypes (37 of 58) were nominally significantly associated with CNV carrier status (p<0·05), all in the direction of increasing risk for developing the phenotype (Table 2 and Figure 1). After Bonferroni correction for 58 phenotypes (p<8·6×10^−4^), 19 of the 37 associations remained significant. Diabetes, cardiovascular, respiratory and renal problems were prominent amongst the most significant results, as was “death during follow-up”. The most significant result was for hypertension, but this was also the most common phenotype, diagnosed in 31.6% of participants, and had a modest OR of 1·28. The three phenotypes with the highest odds ratios were obesity (OR=1·69, f=2·5% of participants), renal failure (OR=1·67%, f=2·1%) and the risk of death during the follow-up period (OR=1·67, f=2·9%).

**Table 2.**
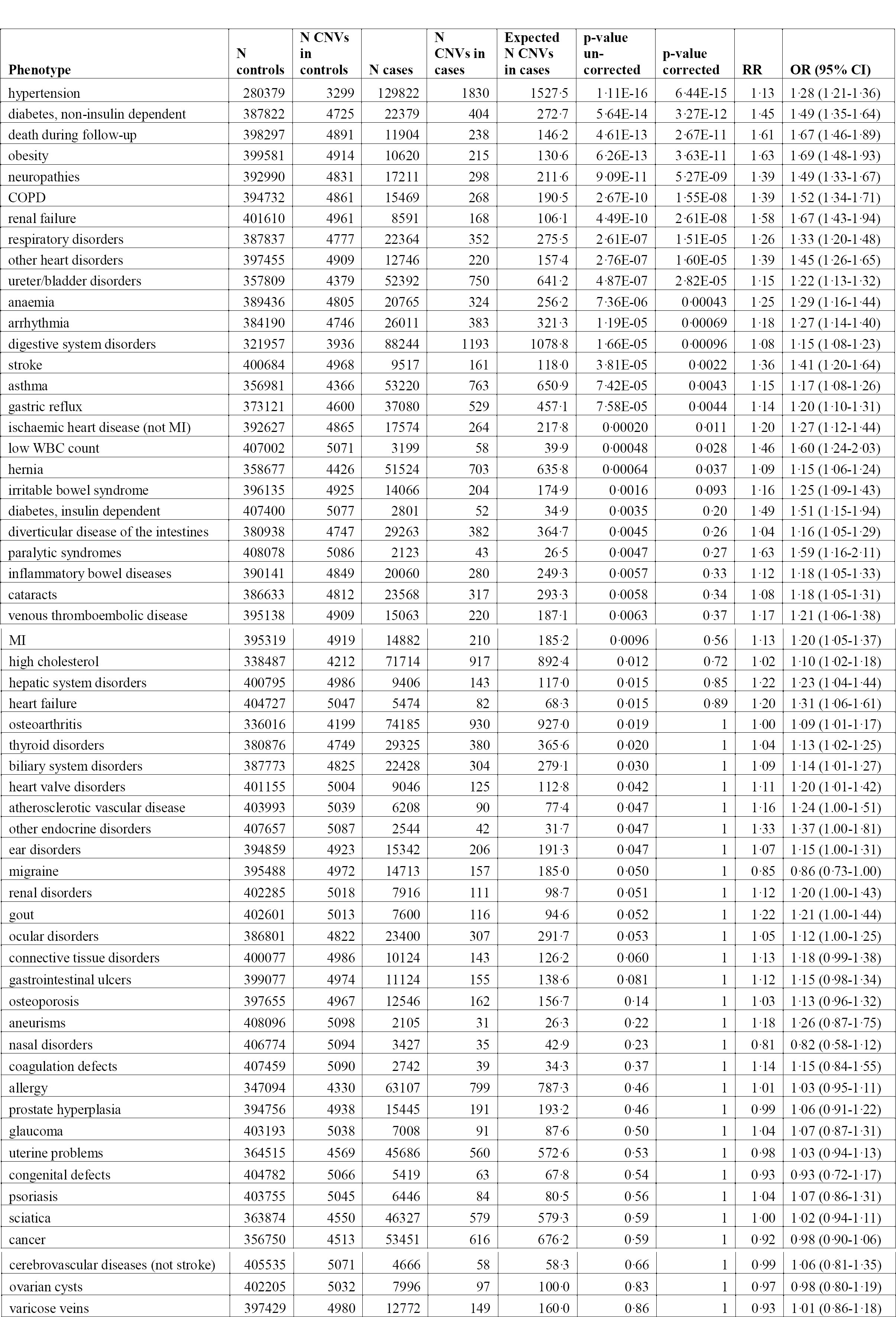
Results for 58 medical phenotypes against the group of 49 rare pathogenic CNVs. “N controls” and “N cases” show the number of people without or with the phenotype. “N CNV controls” and “N CNV cases” refer to the number of CNVs found in these controls and cases. “Expected N CNV cases” is the expected number of CNVs among cases, assuming control frequencies. “OR, 95% CI of OR” and “p-values” refer to the results from Firth logistic regression analysis. “Corrected p-value” is the Bonferroni corrected significance for 58 tests (phenotypes). “RR” stands for the uncorrected risk ratio (relative risk).

| Phenotype | N controls | N CNVs in controls | N cases | N CNVs in cases | Expected N CNVs in cases | p-value un-corrected | p-value corrected | RR | OR (95% CI) |
| --- | --- | --- | --- | --- | --- | --- | --- | --- | --- |
| hypertension | 280379 | 3299 | 129822 | 1830 | 1527.5 | 1.11E-16 | 6.44E-15 | 1.13 | 1.28 (1.21-1.36) |
| diabetes, non-insulin dependent | 387822 | 4725 | 22379 | 404 | 272.7 | 5.64E-14 | 3.27E-12 | 1.45 | 1.49 (1.35-1.64) |
| death during follow-up | 398297 | 4891 | 11904 | 238 | 146.2 | 4.61E-13 | 2.67E-11 | 1.61 | 1.67 (1.46-1.89) |
| obesity | 399581 | 4914 | 10620 | 215 | 130.6 | 6.26E-13 | 3.63E-11 | 1.63 | 1.69 (1.48-1.93) |
| neuropathies | 392990 | 4831 | 17211 | 298 | 211.6 | 9.09E-11 | 5.27E-09 | 1.39 | 1.49 (1.33-1.67) |
| COPD | 394732 | 4861 | 15469 | 268 | 190.5 | 2.67E-10 | 1.55E-08 | 1.39 | 1.52 (1.34-1.71) |
| renal failure | 401610 | 4961 | 8591 | 168 | 106.1 | 4.49E-10 | 2.61E-08 | 1.58 | 1.67 (1.43-1.94) |
| respiratory disorders | 387837 | 4777 | 22364 | 352 | 275.5 | 2.61E-07 | 1.51E-05 | 1.26 | 1.33 (1.20-1.48) |
| other heart disorders | 397455 | 4909 | 12746 | 220 | 157.4 | 2.76E-07 | 1.60E-05 | 1.39 | 1.45 (1.26-1.65) |
| ureter/bladder disorders | 357809 | 4379 | 52392 | 750 | 641.2 | 4.87E-07 | 2.82E-05 | 1.15 | 1.22 (1.13-1.32) |
| anaemia | 389436 | 4805 | 20765 | 324 | 256.2 | 7.36E-06 | 0.00043 | 1.25 | 1.29 (1.16-1.44) |
| arrhythmia | 384190 | 4746 | 26011 | 383 | 321.3 | 1.19E-05 | 0.00069 | 1.18 | 1.27 (1.14-1.40) |
| digestive system disorders | 321957 | 3936 | 88244 | 1193 | 1078.8 | 1.66E-05 | 0.00096 | 1.08 | 1.15 (1.08-1.23) |
| stroke | 400684 | 4968 | 9517 | 161 | 118.0 | 3.81E-05 | 0.0022 | 1.36 | 1.41 (1.20-1.64) |
| asthma | 356981 | 4366 | 53220 | 763 | 650.9 | 7.42E-05 | 0.0043 | 1.15 | 1.17 (1.08-1.26) |
| gastric reflux | 373121 | 4600 | 37080 | 529 | 457.1 | 7.58E-05 | 0.0044 | 1.14 | 1.20 (1.10-1.31) |
| ischaemic heart disease (not MI) | 392627 | 4865 | 17574 | 264 | 217.8 | 0.00020 | 0.011 | 1.20 | 1.27 (1.12-1.44) |
| low WBC count | 407002 | 5071 | 3199 | 58 | 39.9 | 0.00048 | 0.028 | 1.46 | 1.60 (1.24-2.03) |
| hernia | 358677 | 4426 | 51524 | 703 | 635.8 | 0.00064 | 0.037 | 1.09 | 1.15 (1.06-1.24) |
| irritable bowel syndrome | 396135 | 4925 | 14066 | 204 | 174.9 | 0.0016 | 0.093 | 1.16 | 1.25 (1.09-1.43) |
| diabetes, insulin dependent | 407400 | 5077 | 2801 | 52 | 34.9 | 0.0035 | 0.20 | 1.49 | 1.51 (1.15-1.94) |
| diverticular disease of the intestines | 380938 | 4747 | 29263 | 382 | 364.7 | 0.0045 | 0.26 | 1.04 | 1.16 (1.05-1.29) |
| paralytic syndromes | 408078 | 5086 | 2123 | 43 | 26.5 | 0.0047 | 0.27 | 1.63 | 1.59 (1.16-2.11) |
| inflammatory bowel diseases | 390141 | 4849 | 20060 | 280 | 249.3 | 0.0057 | 0.33 | 1.12 | 1.18 (1.05-1.33) |
| cataracts | 386633 | 4812 | 23568 | 317 | 293.3 | 0.0058 | 0.34 | 1.08 | 1.18 (1.05-1.31) |
| venous thromboembolic disease | 395138 | 4909 | 15063 | 220 | 187.1 | 0.0063 | 0.37 | 1.17 | 1.21 (1.06-1.38) |
| MI | 395319 | 4919 | 14882 | 210 | 185.2 | 0.0096 | 0.56 | 1.13 | 1.20 (1.05-1.37) |
| high cholesterol | 338487 | 4212 | 71714 | 917 | 892.4 | 0.012 | 0.72 | 1.02 | 1.10 (1.02-1.18) |
| hepatic system disorders | 400795 | 4986 | 9406 | 143 | 117.0 | 0.015 | 0.85 | 1.22 | 1.23 (1.04-1.44) |
| heart failure | 404727 | 5047 | 5474 | 82 | 68.3 | 0.015 | 0.89 | 1.20 | 1.31 (1.06-1.61) |
| osteoarthritis | 336016 | 4199 | 74185 | 930 | 927.0 | 0.019 | 1 | 1.00 | 1.09 (1.01-1.17) |
| thyroid disorders | 380876 | 4749 | 29325 | 380 | 365.6 | 0.020 | 1 | 1.04 | 1.13 (1.02-1.25) |
| biliary system disorders | 387773 | 4825 | 22428 | 304 | 279.1 | 0.030 | 1 | 1.09 | 1.14 (1.01-1.27) |
| heart valve disorders | 401155 | 5004 | 9046 | 125 | 112.8 | 0.042 | 1 | 1.11 | 1.20 (1.01-1.42) |
| atherosclerotic vascular disease | 403993 | 5039 | 6208 | 90 | 77.4 | 0.047 | 1 | 1.16 | 1.24 (1.00-1.51) |
| other endocrine disorders | 407657 | 5087 | 2544 | 42 | 31.7 | 0.047 | 1 | 1.33 | 1.37 (1.00-1.81) |
| ear disorders | 394859 | 4923 | 15342 | 206 | 191.3 | 0.047 | 1 | 1.07 | 1.15 (1.00-1.31) |
| migraine | 395488 | 4972 | 14713 | 157 | 185.0 | 0.050 | 1 | 0.85 | 0.86 (0.73-1.00) |
| renal disorders | 402285 | 5018 | 7916 | 111 | 98.7 | 0.051 | 1 | 1.12 | 1.20 (1.00-1.43) |
| gout | 402601 | 5013 | 7600 | 116 | 94.6 | 0.052 | 1 | 1.22 | 1.21 (1.00-1.44) |
| ocular disorders | 386801 | 4822 | 23400 | 307 | 291.7 | 0.053 | 1 | 1.05 | 1.12 (1.00-1.25) |
| connective tissue disorders | 400077 | 4986 | 10124 | 143 | 126.2 | 0.060 | 1 | 1.13 | 1.18 (0.99-1.38) |
| gastrointestinal ulcers | 399077 | 4974 | 11124 | 155 | 138.6 | 0.081 | 1 | 1.12 | 1.15 (0.98-1.34) |
| osteoporosis | 397655 | 4967 | 12546 | 162 | 156.7 | 0.14 | 1 | 1.03 | 1.13 (0.96-1.32) |
| aneurisms | 408096 | 5098 | 2105 | 31 | 26.3 | 0.22 | 1 | 1.18 | 1.26 (0.87-1.75) |
| nasal disorders | 406774 | 5094 | 3427 | 35 | 42.9 | 0.23 | 1 | 0.81 | 0.82 (0.58-1.12) |
| coagulation defects | 407459 | 5090 | 2742 | 39 | 34.3 | 0.37 | 1 | 1.14 | 1.15 (0.84-1.55) |
| allergy | 347094 | 4330 | 63107 | 799 | 787.3 | 0.46 | 1 | 1.01 | 1.03 (0.95-1.11) |
| prostate hyperplasia | 394756 | 4938 | 15445 | 191 | 193.2 | 0.46 | 1 | 0.99 | 1.06 (0.91-1.22) |
| glaucoma | 403193 | 5038 | 7008 | 91 | 87.6 | 0.50 | 1 | 1.04 | 1.07 (0.87-1.31) |
| uterine problems | 364515 | 4569 | 45686 | 560 | 572.6 | 0.53 | 1 | 0.98 | 1.03 (0.94-1.13) |
| congenital defects | 404782 | 5066 | 5419 | 63 | 67.8 | 0.54 | 1 | 0.93 | 0.93 (0.72-1.17) |
| psoriasis | 403755 | 5045 | 6446 | 84 | 80.5 | 0.56 | 1 | 1.04 | 1.07 (0.86-1.31) |
| sciatica | 363874 | 4550 | 46327 | 579 | 579.3 | 0.59 | 1 | 1.00 | 1.02 (0.94-1.11) |
| cancer | 356750 | 4513 | 53451 | 616 | 676.2 | 0.59 | 1 | 0.92 | 0.98 (0.90-1.06) |
| cerebrovascular diseases (not stroke) | 405535 | 5071 | 4666 | 58 | 58.3 | 0.66 | 1 | 0.99 | 1.06 (0.81-1.35) |
| ovarian cysts | 402205 | 5032 | 7996 | 97 | 100.0 | 0.83 | 1 | 0.97 | 0.98 (0.80-1.19) |
| varicose veins | 397429 | 4980 | 12772 | 149 | 160.0 | 0.86 | 1 | 0.93 | 1.01 (0.86-1.18) |

**Figure 1:**
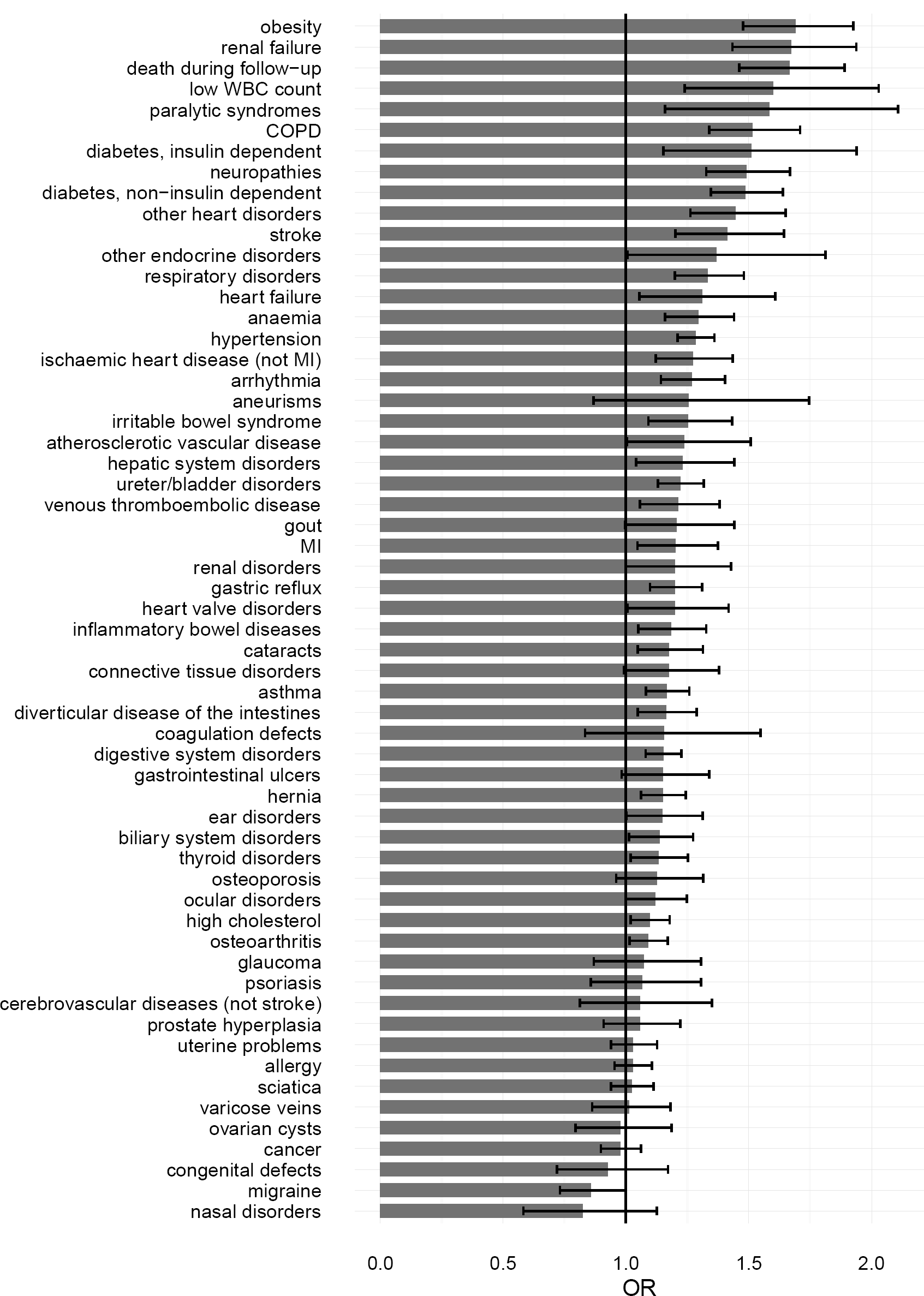
ORs and 95% CI for the ORs for developing each of the 58 tested phenotypes in carriers of any one of the 49 rare pathogenic CNVs. The phenotypes are ordered by the strength of the OR.

### Effects of individual CNVs on phenotypes

Identifying links between specific CNVs and specific disorders is of high clinical importance. The supplementary information (Table S3) presents the effect of each of the 54 CNVs on each of the 58 phenotypes. Figure 2 demonstrates, as an example, the ORs of arguably the most important phenotype, death during follow-up, for carriers of each of the 54 CNVs, ordered by the statistical strength of the association (strongest p-value on top). The horizontal black line demarcates the 17 CNVs that are nominally significantly associated with increased risk for death.

**Figure 2:**
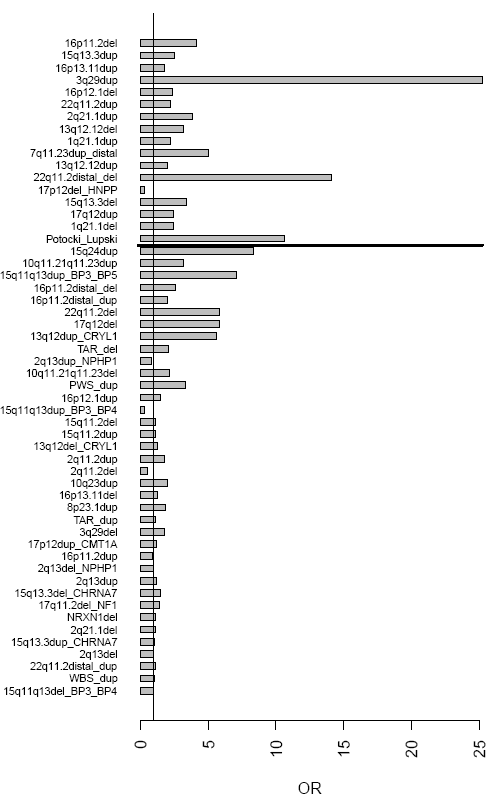
ORs for dying during the follow-up interval, up to 2016, for carriers of each of the 54 CNVs tested in this study. The CNVs are ordered by the strength of the significance (strongest result on top, for 16p11·2 deletions). The horizontal line demarcates the last nominally significant result (p<0·05).

After Bonferroni correction for 3132 tests (a project-wide significant p-value threshold of 1·6×10^−5^), 18 CNV/phenotype comparisons remained significant (Table 3). Using Benjamini-Hochberg FDR=0.1 we report an additional 57 significant associations (Table S4). Clinicians might decide to also consider consequences of CNVs that do not survive multiple testing correction, therefore we present both corrected and uncorrected p-values in Table S3 and the website.

**Table 3.** List of results that survive Bonferroni correction for 3132 tests. Variable names follow the nomenclature for Table 2.

| Phenotype | CNV | N controls | N CNVs controls | N cases | N CNVs cases | Expected N CNVs in cases | RR | p-value | OR(95%CI) |
| --- | --- | --- | --- | --- | --- | --- | --- | --- | --- |
| anaemia | 16p11·2del | 384721 | 90 | 20461 | 20 | 4·8 | 3·6 | 6·94E-08 | 4·8(2·9-7·6) |
| asthma | 16p11·2del | 352692 | 77 | 52490 | 33 | 11·5 | 2·3 | 3·24E-06 | 2·8(1·9-4·2) |
| asthma | 15q13·3del | 352640 | 25 | 52474 | 17 | 3·7 | 3·1 | 5·75E-06 | 4·6(2·5-8·5) |
| diabetes insulin dependent | 17q12del | 402328 | 5 | 2753 | 4 | 0·0 | 65·5 | 2·50E-08 | 121·5(31·9-440·3) |
| diabetes non-insulin dependent | 16p11·2del | 383179 | 82 | 22003 | 28 | 4·7 | 4·7 | 1·49E-14 | 7·9(5·0-12·1) |
| diabetes non-insulin dependent | 16p11·2distal_del | 383139 | 42 | 21991 | 16 | 2·4 | 5·1 | 6·22E-09 | 7·9(4·2-14·0) |
| gastric reflux | 22q11·2dup | 368750 | 229 | 36602 | 51 | 22·7 | 2·0 | 1·36E-07 | 2·5(1·8-3·3) |
| hypertension | 16p11·2del | 277136 | 56 | 128046 | 54 | 25·9 | 1·6 | 7·04E-08 | 3·0(2·0-4·5) |
| hypertension | 16p12·1del | 277213 | 133 | 128105 | 113 | 61·5 | 1·4 | 8·16E-08 | 2·1(1·6-2·7) |
| hypertension | 16p13·11dup | 277590 | 510 | 128310 | 318 | 235·7 | 1·2 | 2·28E-06 | 1·4(1·2-1·7) |
| neuropathies | 17p12dup_CMT1A | 388224 | 65 | 16972 | 59 | 2·8 | 11·4 | 4·70E-47 | 22·1(15·5-31·5) |
| neuropathies | 17p12del_HNPP | 388370 | 211 | 16939 | 26 | 9·2 | 2·6 | 8·08E-06 | 2·9(1·9-4·2) |
| obesity | 16p11·2del | 394758 | 91 | 10424 | 19 | 2·4 | 6·7 | 7·58E-12 | 8·7(5·2-13·9) |
| obesity | 16p12·1del | 394890 | 223 | 10428 | 23 | 5·9 | 3·6 | 7·55E-08 | 4·1(2·6-6·1) |
| obesity | 16p11·2distal_del | 394716 | 49 | 10414 | 9 | 1·3 | 6·0 | 4·64E-06 | 7·7(3·6-14·8) |
| osteoarthritis | 16p11·2del | 331895 | 78 | 73287 | 32 | 17·2 | 1·6 | 1·22E-05 | 2·8(1·8-4·2) |
| renal failure | 16p11·2del | 396748 | 99 | 8434 | 11 | 2·1 | 4·8 | 7·56E-07 | 7·3(3·7-13·1) |
| renal failure | 16p12·1del | 396878 | 229 | 8440 | 17 | 4·9 | 3·3 | 6·55E-06 | 3·9(2·3-6·1) |

### Homozygous deletions

We examined the data for homozygous deletions. Not surprisingly for rare pathogenic CNVs and relatively healthy middle or old-aged people, only four such instances were found. Three of these clustered in a single locus, at 2q13 (110,21-110,34Mb), affecting the gene NPHP1. Homozygous deletions at this locus are known to cause the kidney disorder juvenile nephronophthisis. All three Biobank individuals with homozygous deletions at 2q13 had renal failure (Fisher exact test p=9·2×10^−6^).

## Discussion

With half a million participants, long duration of follow-up and unbiased gathering of phenotypic information, the UK Biobank provides a unique resource to test the effect of genetic variation on multiple phenotypes in middle and old age. We show that the microarray data is of a high quality, with only ~5% of arrays not surviving standard QC filtering, and provides reliable calls for the vast majority of known pathogenic CNVs. The frequencies of these CNVs are extremely similar to those among controls in other studies, as we reported previously.^9^ If our methods can detect true associations, we would expect that the best-established medical consequences of CNVs would be among our top hits. Indeed, the 18 findings that survive conservative Bonferroni correction (Table 3) contain some well-known associations, thus providing the best proof of principle that these methods will detect true associations. These include the expected associations with neuropathies and 17p12 deletions and duplications;^14^ asthma and 16p11 −2 deletions;^15^ obesity and 16p11·2 deletions and 16p11·2 distal deletions;^16,17^ diabetes and deletions at 17q12 (also called “renal cysts and diabetes syndrome”).^18^ We now provide data showing that in adults some of these effects lead to a high incidence of non-insulin dependent diabetes mellitus, osteoarthritis and hypertension.

The set of 49 CNVs (after excluding the five relatively common ones) increased risk for more than half of the tested medical phenotypes (Table 2 and Figure 1). It follows that many individual CNVs increase risk for specific phenotypes. Indeed, we observed 382 nominally significant CNV/phenotype associations, instead of the expected 156 (Figure S1), with 75 of those significant at a B-H FDR=0·1 (Table S4) and 18 surviving conservative Bonferroni correction for all the tests performed in this project (Table 3). The five relatively common CNVs produced only one significant result at B-H FDR=0.1, despite having much higher statistical power to detect associations with similar ORs, due to their higher frequencies. This is in line with the notion that stronger selection pressure lowers the CNV frequencies.^19^

The CNV impacting the highest number of phenotypes is the deletion at 16p11·2, with 28 nominally significantly associated phenotypes (15 of them significant at B-H FDR=0·1), Table S4/16p11·2del). Apart from the phenotypes that are expected comorbidities of the recognised high body mass index BMI (hypertension, diabetes, osteoarthritis, heart failure),^16^ there were others that are not obviously linked to a high BMI, for example COPD, cataract, psoriasis, anaemia and renal problems. We should point out that this does not make the 16p11·2 deletion the most pathogenic CNV, as significance depends also on CNV frequency, which is low for the highly pathogenic CNVs. The most pathogenic CNVs are underrepresented in the Biobank, as the participants are middle-aged and participation is subject to “healthy volunteer” selection bias.^20^ For example there were only 10 carriers of 22q11·2 deletions, instead of the expected ~100 (the rate of this deletion among newborns is ~1:4,000).^7^

The increased risk for medical morbidities observed in CNV carriers is unlikely to be due to the presence of early neurodevelopmental disorders or schizophrenia in carriers, as the UK Biobank population has largely escaped such conditions: only 35 of the 16,167 people who had one of the tested CNVs had schizophrenia and only four declared a diagnosis of autism. We did not control for cognitive ability, education, smoking, area of living, household income and other socio-economic factors. Although these have established effects on health outcomes, they may also be influenced by the CNV carrier status. Indeed, we have previously shown that these CNVs affect people’s educational and occupational attainment.^9^

For most phenotypes, the differences between the expected and observed CNVs in cases (Table 2) cannot be accounted for by the significant individual CNV associations alone. As an extreme example, the phenotype arrhythmia is highly statistically associated with CNV carrier status, but has no individual CNV association, even at a nominal level of significance. This suggests that more associations are true but could not reach statistical significance due to the small number of observations.

Our results can guide clinicians into monitoring carriers of specific CNVs. For example, it appears that people with 16p11·2 duplications (who tend to be tall and slim) should be monitored for osteoporosis, renal failure and venous thromboembolic disease and could benefit from treatment of irritable bowel syndrome, as it might contribute towards poor food intake. Other examples are carriers of 16p12·1 deletions, who may benefit from monitoring for anaemia, COPD, uterine and ovarian problems, while those with 1q21·1 deletions might need regular eye checks. Although these CNVs are individually rare, as a group they are found in over 1% of adults in the general population (after excluding the five relatively common ones), making them important sources of morbidity and mortality.

CNVs are likely to have more specific disease outcomes, rather than leading to the broad disease groups that we defined. While any association can be tested, it is more appropriate for such detailed analysis to be conducted by researchers interested in specific phenotypes, and to be hypothesis-driven, rather than *en masse* against all possible outcomes.

Finally, our lists of phenotypes associated with CNVs should provide researchers with another avenue for the elucidation of pathophysiological disease mechanisms.

## Acknowledgements

This research has been conducted using the UK Biobank Resource under Application Number 14421. The work at Cardiff University was funded by the Medical Research Council (MRC) Centre Grant (MR/L010305/1) and Program Grant (G0800509). KC and MBS received PhD studentships funded by the MRC UK. KMK is funded by a Research Training Fellowship from the Wellcome Trust, London.

